# Loss of Fgf-responsive Pea3 transcription factors results in ciliopathy-associated phenotypes during early zebrafish development

**DOI:** 10.1101/2020.09.04.283804

**Authors:** Matt E. McFaul, Neta Hart, Bruce W. Draper

## Abstract

FGF signaling is used reiteratively during development and elicits several different responses, such as cell proliferation, differentiation, or migration. We parsed the complex FGF intracellular response by creating triple homozygous mutants in the Pea3 subgroup of ETS transcription factors, designated *3etv* mutants. The Pea3 proteins Etv4 and Etv5 are expressed in areas of FGF activity; however, their role in FGF signal transduction as either positive or negative modulators of FGF activity was unclear. Using *3etv* mutants, we found these genes act redundantly and have phenotypes consistent with known FGF defects in inner ear, pectoral fin, and posterior mesoderm development. Additionally, we uncovered a novel role for the FGF/Pea3 pathway during body axis straightening. *3etv* larvae develop a curly-tail up (CTU) phenotype that we linked to mis-regulation of the polycystin and urotensin pathways, which have opposing actions to ensure a straight body orientation along the dorsal-ventral axis. We find that the Etv4/5 transcription factors act as positive regulators of FGF signaling and propose a model where Etv4/5 are required for cilia function downstream of Fgf8a.

**Summary Statement:** Pea3 transcription factor triple mutants reveal a role for FGF signaling in balancing polycystin and urotensin signaling to achieve a straight body axis.

## Introduction

The fibroblast growth factor (FGF) signaling pathway is a critical cellular communication mechanism used repeatedly during embryonic development. FGF signaling occurs in several different contexts – acting as an extracellular short range, distant endocrine, or intracellular signal – and in turn can activate intracellular response cascades such as the mitogen-activated protein kinase (MAPK), PI3K-AKT, Jun/Fos and PLC-ɣ pathways (Geary and Labonne, 2018; Wang et al., 1994; Yang et al., 2009). The vast array of signaling paradigms and cellular responses allows FGF to regulate diverse processes such as cell proliferation, migration, and organ morphogenesis and patterning in different tissues throughout development and during adult homeostasis. The role of FGF signaling in development has been well studied at the level of FGF ligands and FGF receptors (FGFR); however, less is known about the role of downstream transcription factor mediators of FGF signaling.

The E26-transformation specific (ETS) protein family is comprised of 28 mammalian transcription factors characterized by a conserved winged helix-loop-helix ETS DNA-binding domain and are regulated by the MAPK pathway. (Hollenhorst et al., 2011; Wasylyk et al., 1998). Expression and transcriptional activity of the Polyoma enhancer activator Pea3 subclass of ETS transcription factors are regulated by FGF signaling via the Erk and Jnk branches of the MAPK cascade (O’Hagan et al., 1996). Because Pea3 genes are routinely expressed in response to FGF signaling in many different developmental contexts, they are considered direct targets and general downstream mediators of FGF activity (Brent and Tabin, 2004; Raible and Brand, 2001; Roehl and Nüsslein-Volhard, 2001). Despite these observations, the role of Pea3 transcription factors and whether they are necessary for regulating FGF output during development is not well characterized, with conflicting reports in the literature. For example, Pea3 genes can directly induce negative feedback of FGF signaling and therefore attenuate the effects of FGF, while other studies find Pea3 gene mutants have defects consistent with loss of FGF suggesting a positive role in signal mediation (Garg et al., 2018; Garg et al., 2020; Herriges et al., 2015; Znosko et al., 2010). The closely related ETS-domain containing proteins, Etv1, Etv4, and Etv5 make up the Pea3 family of transcription factors. Though they share sequence homology, there is considerable evidence in several organism that Etv1 takes on different roles than Etv4 and Etv5 (Faedo et al., 2010; Herriges et al., 2015; Roussigné and Blader, 2006; Zhang et al., 2009). In mice, Etv4 and Etv5 share similar expression patterns and mutational analyses have found they are functionally redundant in many developmental events, for example in the eye lens, dorsal root ganglion cells, and limb formation (Fontanet et al., 2013; Garg et al., 2020; Lettice et al., 2012).

The roles of Etv4/5 transcription factors have previously been investigated in zebrafish (*Danio rerio*) embryonic development using morpholino oligo (MO) knockdown and it was found that, similar to mice, these genes appear to function redundantly, as phenotypes are only observed only when the *etv4* and *etv5* genes are simultaneously targeted (Mao et al., 2009; Zhang et al., 2009; Znosko et al., 2010). The zebrafish genome contains a single homolog of *Etv4*, called *etv4* (formerly *pea3*) and two ohnologs of *Etv5*, called *etv5a* and *etv5b* (formerly *etv5* and *erm*, respectively); the two Etv5 genes are a result of the teleost-specific whole genome duplication event (Chen et al., 2013; Liu and Patient, 2008). In zebrafish *etv4, etv5a*, and *etv5b*, but not *etv1*, function downstream of FGF signaling as they are expressed in similar domains to developmentally important FGF ligands and their expression is significantly reduced or lost when FGF signaling is inhibited (Roussigné and Blader, 2006). Triple morphant *etv4;etv5a;etv5b* knockdown zebrafish embryos have defects in left-right (LR) patterning, midbrain-hindbrain boundary (MHB), cardiac, and limb development. However, MO are only effective for gene knockdown during very early development, generally before 48 hours post fertilization (hpf), so it remains to be determined what roles these genes may play during later development.

Here we describe the generation and characterization of genetic mutations in *etv4, etv5a*, and *etv5b*. We find that these three genes function redundantly as all single and double mutant fish are viable and fertile, in accordance with previous studies (Garg et al., 2020; Znosko et al., 2010). Triple homozygous mutants are inviable and have many developmental defects consistent with reductions in FGF signaling. Additionally, and not consistent with known roles of FGF signaling, we find that triple mutants have tail curvature defects that develop by 2 days postfertilization (dpf) and are remarkably similar to those caused by mutations that disrupt the function of polycystin genes *pkd1* and *pkd2*, which are thought to function in primary cilia mechanosensation. Finally, we confirm that this defect is due to reductions in FGF signaling output and arises from dysregulation of polycystin and urotensin pathways. We conclude that the Etv4/5 transcription factors act as positive regulators of FGF and based on the many defects observed in triple mutants, we propose a model in which Pea3 activity is required for cilia function downstream of Fgf8a. This suggests the FGF/MAPK/Pea3 signaling axis, in addition to the FGF/Ca^2+^ axis, is required for cilia activity (Schneider et al., 2019).

## Results

### Redundancy and genetic compensation in Pea3 transcription factors

FGF signaling is known to play key roles during multiple stages of development, and cells responding to FGF signaling express one or more of the *etv4, etv5a* and/or *etv5b* genes (Brent and Tabin, 2004; Raible and Brand, 2001; Roehl and Nüsslein-Volhard, 2001). To identify the specific developmental processes that Pea3 factors regulate, we used CRISPR/Cas9 gene editing to produce frameshift mutations in the zebrafish *etv4, etv5a* and *etv5b* genes. Protein structures of the Pea3 transcription factors include a conserved N-terminal activation domain and a C-terminal ETS DNA-binding domain, each flanked by auto-inhibitory sequences that regulate transcriptional activity (Fig. 1A; Bojović and Hassell, 2001; Currie et al., 2017; Greenall et al., 2001). To ensure non-functional alleles were produced, we designed Cas9 single-guide RNAs (sgRNA) to target DNA double-strand break formation upstream of the ETS DNA binding domain in all three genes. We identified insertion/deletion (indel) mutations in the coding regions of each gene that resulted in frameshifts prior to the DNA binding domain and confirmed the mutant alleles with genomic DNA and cDNA Sanger sequencing. (Fig. 1A; Fig. S1).

**Figure 1.**
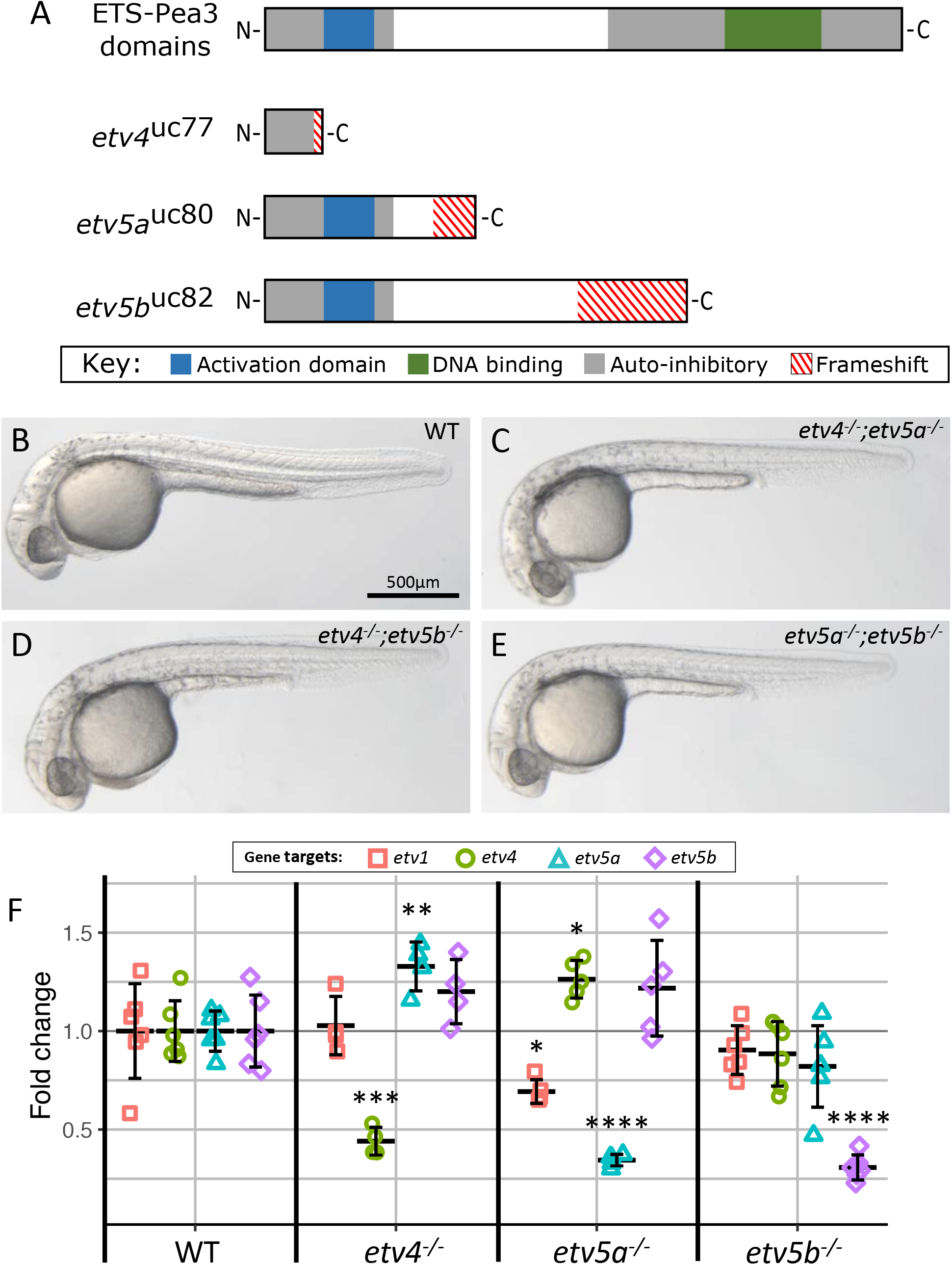
Redundancy and genetic compensation in Pea3 transcription factors. (A) Schematic of Pea3 transcription factors protein domains, along with truncations in *etv4, etv5a*, and *etv5b* mutant proteins in CRISPR/Cas9 induced frameshift mutations. (B-E) Light microscope images of wild-type and Pea3 double mutant combinations at 30 hpf. (F) Fold change values of Pea3 transcription factors measured by RT-qPCR for etv4^-/-^, etv5a^-/-^, and etv5b^-/-^ single mutants relative to wild-type in 24 hpf embryos. Individual fold change values for biological replicates are plotted with the mean fold change and standard deviation error bars.

We generated homozygous single mutants and all three combinations of double mutants and found that in each case the mutant animals were viable and fertile. Figure 1B-D shows representative wild-type and double mutant embryos at 30 hpf. These results are consistent with previously reported analysis using MO knockdowns (Mao et al., 2009; Zhang et al., 2009; Znosko et al., 2010). In addition to zygotic expression, *etv4, etv5a* and *etv5b* are all expressed maternally (Chen et al., 2013; Raible and Brand, 2001; Roussigné and Blader, 2006). To determine if maternal expression of these genes is important for early development, we produced maternal-zygotic (MZ) single and double mutant combinations for *etv4, etv5a*, and *etv5b* that lacked both maternal and zygotic gene products. We found that all MZ double mutant combinations were viable and fertile as adults.

It has recently been discovered that mutations resulting in a premature stop codon in one gene can lead to the transcriptional upregulation of another related gene by a mechanism termed genetic compensation (El-Brolosy et al., 2019). Genetic compensation appears to be triggered by the degradation of the mutant transcript by nonsense-mediated decay (NMD). This mechanism likely explains why in some cases genetic mutations engineered by gene editing can result in weaker phenotypes than those produced using translation-blocking MOs, as the latter does not cause NMD (El-Brolosy et al., 2019; Ma et al., 2019; Rossi et al., 2015). To determine if genetic compensation is triggered by our frameshift alleles, we performed RT-qPCR to compare the transcript levels of the four Pea3 genes *etv1, etv4, etv5a*, and *etv5b* in single mutant embryos for *etv4, etv5a* or *etv5b*. In all three mutants we observed significant reduction in the level of the mutant transcript relative to wild-type controls, which is a hallmark of NMD (Fig. 1F). Furthermore, we found evidence for a compensatory relationship between *etv4* and *etv5a*, where the expression of *etv5a* is upregulated in *etv4* mutants and vice versa. However, it is unlikely this increased expression has a rescuing effect as *etv4;etv5a* double mutants are also viable and fertile. Additionally, we found no increased expression of *etv4* or *etv5a* in *etv5b* mutants. Finally, transcript levels for the fourth Pea3 family member, *etv1*, were not altered in *etv4* or *etv5b* mutants relative to wild-type but were significantly reduced in *etv5a* mutants. Taken together our results suggest that genetic compensation is unlikely to explain why neither single or double mutants have developmental defects, but instead argues for genetic redundancy.

### *etv4;etv5a;etv5b* triple mutants have defects consistent with reduced FGF signaling

To more directly test for functional redundancy, we produced *etv4;etv5a;etv5b* triple mutants, hereafter referred to *3etv* mutants. To our surprise, *3etv* mutant embryos at 30 hpf looked grossly normal when compared to wild-type siblings; however, upon closer examination we noticed several important differences (Fig. 2A, B). First, at 30 hpf *3etv* mutants have shorter body length than wild-type (Fig. 2A, B, C). Second, the otic vesicles of *3etv* mutants were small and 50% contained only a single otolith, while wild-type embryos had two otoliths in each otic vesicle as expected (Fig. 2A’, B’, D). Third, whereas somite boundaries in wild-type embryos are characteristically chevron-shaped with readily apparent boundaries, those in mutant embryos were U-shaped with less well-defined boundaries (Fig. 2A’’, B’’). In fact, it is possible to identify mutant embryos as early as 10 hpf based solely on these somite boundary defects (Fig. 2E, E’, F, F’). Fourth, at 3 dpf mutant embryos have pectoral fin development and outgrowth defects as evident by shortened and misshapen fins that lack actinotrichia (Fig. 2G-J). Fifth, we identified craniofacial defects in *3etv* mutants. In the viscerocrania, mutants had a shortened Meckel’s cartilage and abnormal formation of the hyomandibular cartilage (Fig. 2K,L). Additionally, they had with incomplete ethmoid plate fusion in the neurocranium (Fig. 2M-P). Finally, in contrast to wild-type embryos that inflate their swim bladders at 4 dpf, the swim bladders of *3etv* mutants never inflate and they do not survive past 13 dpf. All defects so far described for the *3etv* mutants are similar to defects caused by mutations of particular FGF ligands; primarily with loss of Fgf8a activity, though defects associated with Fgf3 and Fgf24 are also present (Draper et al., 2003; Fischer et al., 2003; Gebuijs et al., 2019; Léger and Brand, 2002; Reifers et al., 1998; Trumpp et al., 1999). This is consistent with the proposed role of *etv4, etv5a* and *etv5b* as positive mediators of the FGF signaling pathway.

**Figure 2.**
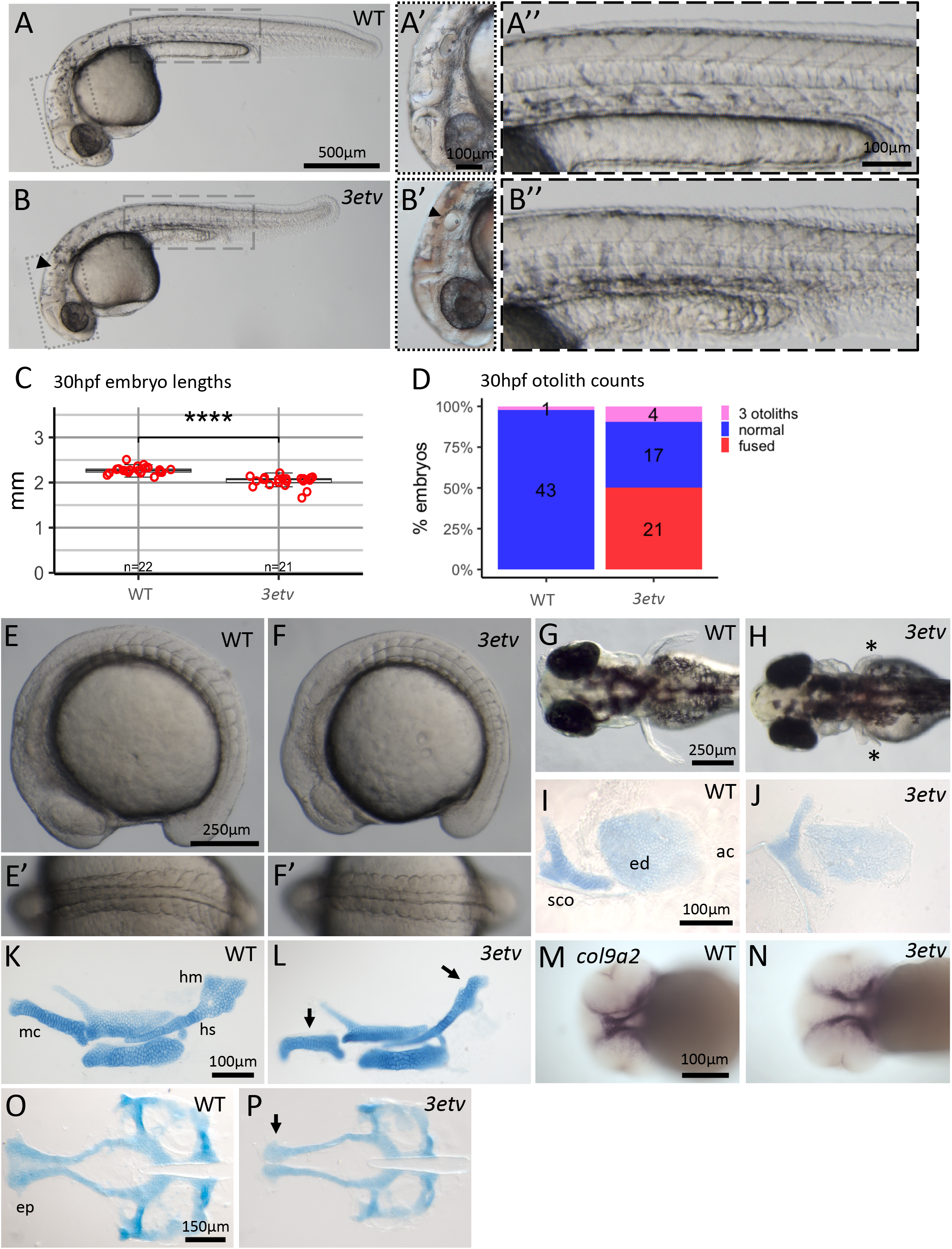
*3etv* mutant phenotypes resemble Fgf defects. 30 hpf light microscope images of wild-type (A-A’’) and *3etv* (B-B’’) embryos. Triple mutants have a reduced body axis length, quantified in (C), and inner ear defects (B’) evidenced by abnormal otolith formation (arrowhead), quantified in (D). (E-F) Lateral view and dorsal view (E’-F’) of wild-type and *3etv* 10 ss embryos. *3etv* mutants can be identified as early as 10 ss by defects in somite boundaries and somite shape. (G-H) Dorsal view of 3 dpf larvae; *3etv* mutants have abnormal, shortened pectoral fins (asterisks) compared to wild-type. (I-J) Alcian blue staining of 6 dpf larvae shoulder girdle and pectoral fin. Mutants form a **sco** but have a misshapen **ed** and lack **ac**. (K-L) Alcian blue staining of viscerocranial cartilage flat-mounted for 6 dpf larvae labeled, *3etv* mutants have defective **mc** and **hm** (arrows). (M-N) Ventral view of *col9a2* ISH staining in 52 hpf embryos labeling developing ethmoid cartilage. (O-P) Alcian blue staining of neurocranial cartilage in 6 dpf larvae. *3etv* mutants have incomplete fusion at the **ep** (arrow). Label abbreviations: scapulocoricoid (sco), endoskeletal disk (ed), actinotrichia (ac), Meckel’s cartilage (mc), hyosymplectic (hs), and hyomandibula (hm), ethmoid plate (ep).

### *3etv* mutants have defects in somite formation

One of the major roles for FGF signaling during early development is to promote the formation and subsequent patterning of posterior mesoderm, and our preliminary analysis suggest that *3etv* mutants have defects in posterior mesoderm development (Amaya et al., 1991; Ciruna et al., 2002; Draper et al., 2003; Griffin et al., 1995; Leerberg et al., 2019; Yamaguchi et al., 1994). To explore this defect, we first compared the expression of the respective axial and paraxial mesoderm markers *tbxta* (formerly *ntl*) and *myod1*, in 11 and 15 somite-stage (ss) embryos (Schulte-Merker et al., 1992; Weinberg et al., 1996). In addition, we co-stained for *pax2a*, a gene expressed in the MHB, early otic vesicle, and in the developing pronephric ducts (Krauss et al., 1991). At 11 ss, *3etv* mutant embryos have a reduction in the overall length of the trunk and tail despite having the same number of somites as wild-type (Fig. 3A,B). The width of the posterior body in 11 ss mutants also appeared to be increased relative to wild-type embryos, which would suggest that the tail length differences are not due to overall developmental delay in mutants, but rather to a defect or delay in convergence and extension of paraxial mesoderm. At 15 ss the overall length of the posterior body is similarly reduced in mutants relative to wild-type, but now there is little to no apparent difference in width (Fig. 3C,D). Analysis of *myod1* expression revealed that the boundaries between somites is much less distinct in mutant relative to wild-type embryos (Fig. 3D). Finally, we also noticed reduction of *pax2a* expression in the MHB and otic placode at 15 ss (Fig. 2C,D), a result consistent with the severe inner ear defects that arise later in *3etv* mutants at 30 hpf (Fig. 2A’, B’). However, despite the MHB having reduced expression of *pax2a* at 15 ss, its development appeared largely normal morphologically in *3etv* mutants at 30 hpf, though we cannot rule out subtle defects (Fig. 2 A’, B’).

**Figure 3.**
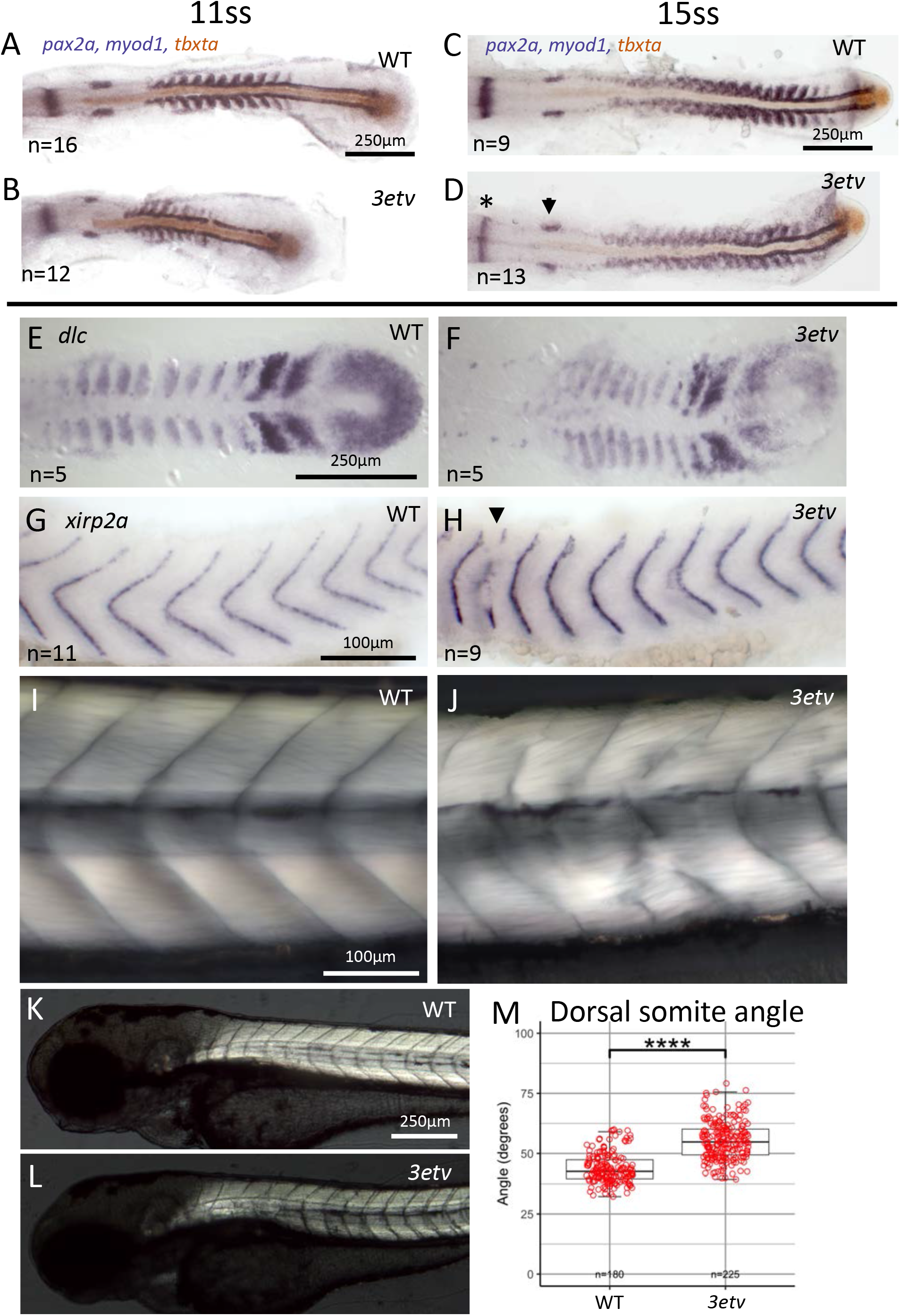
Abnormal somite patterning and boundary defects in *3etv* mutants. (A-D) ISH of *pax2a, myod1*, and *tbxta* in wild-type and *3etv* mutants during somitogenesis. (A-B) At 11 ss *3etv* mutants have reduced *myod1* expression in somites. Expression domains of myod1 in somites are smaller and irregularly shaped in *3etv* mutants compared to wild-type. (C-D) 15 ss embryos continue to have weaker expression of somitic *myod1* and have indistinct boundaries between more posterior somites, conversely there are clearly distinct boundaries between *myod1* domains of adjacent posterior somites in wild-type embryos. Additionally, *3etv* mutants (D) have reduced *pax2a* expression in the MHB (asterisk) and otic vesicle (arrowhead). (E-F) ISH for *dlc* in wild-type and *3etv* embryos at 10 ss. *3etv* embryos have smaller *dlc* expression domains in individual somite boundary segments and somites are compressed into a smaller area (F). (G-H) ISH for *xirp2a* in wild-type and *3etv* embryos at 30 hpf. (G) Wild-type embryos have continuous expression of *xirp2a* in the boundary between adjacent somites in a chevron shaped pattern. (H) Expression of *xirp2a* in *3etv* embryos is U-shaped with gaps in expression in places for some somite boundaries (arrowhead). (I-J) DIC-Nemarski micrographs of wild-type and *3etv* somites in 6 dpf larvae. (J) *3etv* somite boundaries are discontinuous and adjacent somites are fused in places. (K-L) Nemarski-DIC micrographs of wild-type and *3etv* somites and somite boundaries at 6 dpf. (M) Bar graph of curvature angles measured for the dorsal somite boundaries. The first 15 somite angles were measured from at least 12 larvae.

We sought to further characterize the posterior mesoderm defects in *3etv* mutants and investigated somite boundary formation by examining the expression of *deltaC (dlc*). The Delta-Notch ligand *dlc is* known to regulate somite boundary formation and has oscillating expression in presomitic mesoderm cells, but stabilized expression in the posterior half of somites that forms a striped pattern with characteristic spacing between somites (Van Eeden et al., 1996). We found that *3etv* mutants have nearly normal *dlc* expression throughout the paraxial mesoderm, however spacing between the somitic domains is reduced and the expression extends further laterally (Fig. 3E,F). We next examined the expression of *xirp2a*, which is expressed later specifically in cells that localize to the somite boundaries. Analysis of *xirp2a* expression at 30 hpf revealed wild-type embryos had characteristic chevron shaped boundaries and *3etv* mutants formed U-shaped boundaries, as observed above (Fig. 3G,H). In addition, we found several instances in mutant, but not wild-type, embryos where somite boundary formation was incomplete, as evidenced by gaps in *xirp2a* expression (Fig. 3H). At 4 dpf we used polarized light microscopy to observe somite boundaries and found that *3etv* mutants had incomplete myosepta that resulted in the fusion of adjacent somites (Fig. 3I,J). U-shaped somites were still present at 6 dpf in *3etv* mutants by (Fig. 3K,L). Finally, we quantified the shape of the somite boundaries by measuring the angle formed where the dorsal somite boundary intersects with the notochord sheath and found the *3etv* mutants have a significantly wider somite angle than wild-type (Fig. 3M). From these data we conclude that 3Etv transcription factors normally function to regulate development of the posterior mesoderm, including somite boundary formation, consistent with known roles of FGF signaling in this process (Dubrulle et al., 2001; Kawamura et al., 2005; Sawada et al., 2001).

### Transcriptomic characterization of *3etv* mutants

*3etv* embryos have severe somite segmentation and maturation defects. To further profile the *3etv* mutants and determine the genetic networks disrupted in these fish we performed RNA-sequencing for three biological replicates in WT and *3etv* at 10 ss and 30 hpf. Mutants were identified based on developmental phenotypes at both timepoints. As part of the sequencing data quality assessment we performed principal component analysis on log-normalized gene counts and found that all biological replicates clustered tightly according to their age and genotype, confirming the *3etv* mutants were correctly phenotyped and the biological replicates are comparable to one another (Fig. S2). Differential gene expression analysis found 1278 upregulated and 1137 downregulated genes at 10 ss, and 923 upregulated and 1175 downregulated genes at 30 hpf (Tables S3 and S4). These differentially expressed genes (DEG) were determined with an absolute fold change greater than 1.5 and a p-value less than 0.01 (Fig. S2). We performed gene ontology (GO) analysis for the downregulated genes at each timepoint to gain insight into the biological processes activated by the Pea3 transcription factors. Consistent with the defects seen in *3etv* mutants, skeletal muscle development was among the top enriched GO terms at both 10 ss and 30 hpf (Fig 4A,C). We then examined specific gene pathways and cell markers using the RNA-seq data to better understand the somite defects that arise at each timepoint. For the 10 ss dataset we looked at the expression of genes necessary for presomitic mesoderm formation and segmentation, including oscillatory clock genes dlc, *her7*, and *her12*, and found they were downregulated (Fig. 4B). Transcription factors *tbx6, ripply1*, and *ripply2* are critical for the formation of distinct somites; in the absence of these factors *xirp2a* expression is severely disrupted at somite boundaries and adjacent somites are fused together (Ban et al., 2019; Kinoshita et al., 2018). To see if the somite fusion in *3etv* mutants could be attributed to reduced expression of *tbx6* or *ripply1/2* genes we looked at their expression in our 10 ss RNA-seq dataset and found that these genes were also reduced in *3etv* mutants compared to wild-type (Fig. 4B). For the 30 hpf RNA-seq dataset we looked at markers of different somite derivatives: skeletal muscle, myosepta, and sclerotome (Fig. 4D). We found that most skeletal muscle and myosepta markers were downregulated at 30 hpf, whereas sclerotome marker genes were unchanged or slightly upregulated. In all, our data strongly support the conclusion that Etv4, Etv5a and Etv5b function to regulate somite formation and differentiation of somite derivatives.

**Figure 4.**
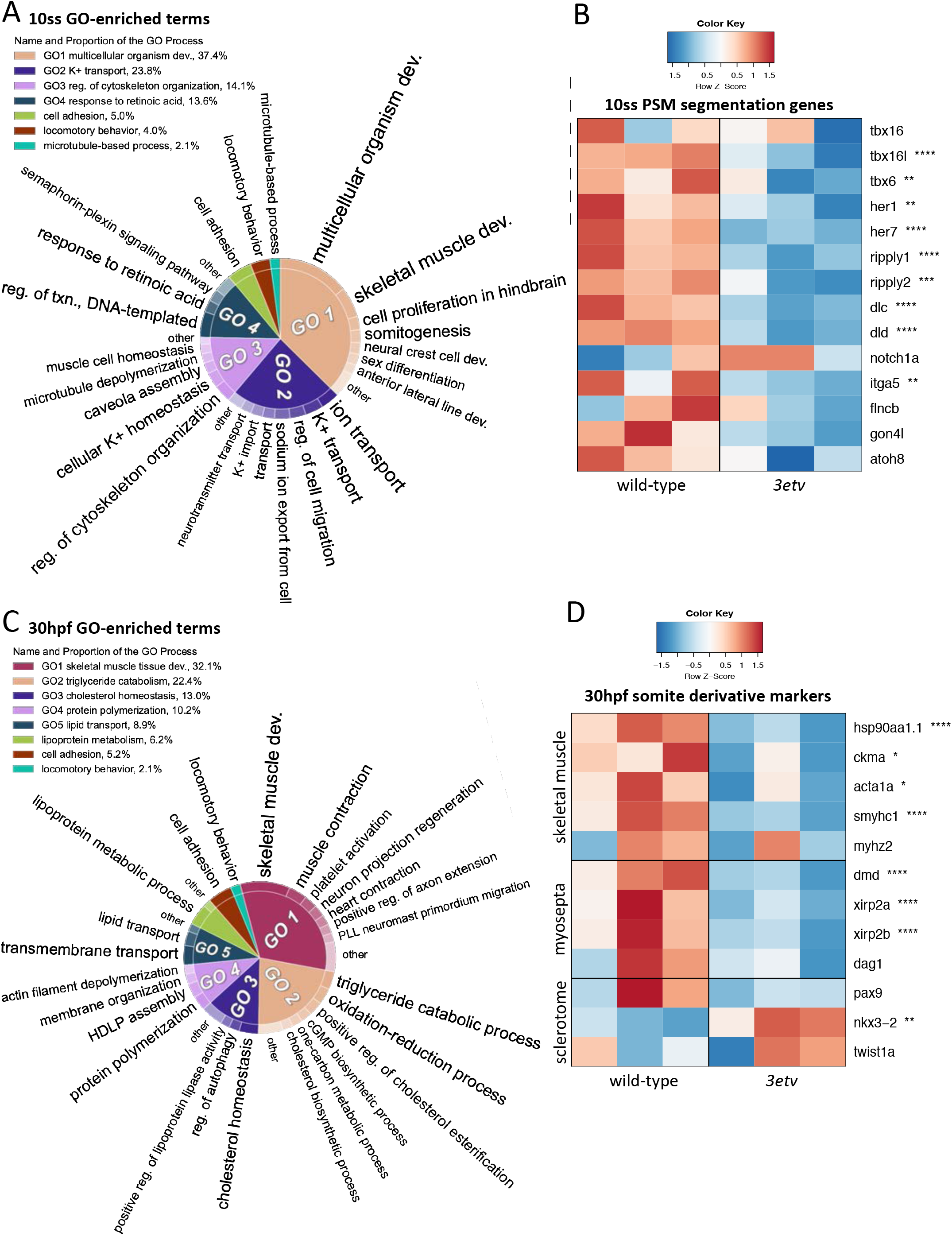
RNA-seq analysis of *3etv* mutants. (A,C) Graphical representation of GO enriched biological processes for significantly downregulated genes at 10 ss and 30 hpf. (B,D) Heatmap representation of RNA-seq gene expression levels for biological replicates in wild-type and *3etv* datasets. Z-score of genes expression for each sample are shown, with reduced expression in blue and increased expression in red. (B) For the 10 ss dataset, genes required for segmentation of the presomitic mesoderm (PSM) and somite boundary formation were measured. (D) At 30 hpf we depicted the expression of various somite derivative marker genes.

### FGF signaling modulates body axis straightening during late embryogenesis

Unexpectedly, *3etv* mutants displayed a striking body axis defect with dorsally curved tails after 2 dpf (Fig. 5A-C). During normal development the trunk and tail initially form with a ventral-ward curvature that then straightens until about 30 hpf when a horizontal body axis is achieved; this process appears to be driven by contractions of muscles located in the dorsal half of each somite (Zhang et al., 2018). This curly-tail up (CTU) phenotype in *3etv* mutants suggests they lose negative feedback to attenuate the tail straightening mechanism, which was a surprising observation as previous studies perturbing FGF signaling have not reported this phenotype. We speculated that the dorsal tail curvature in *3etv* mutants could be independent of FGF signaling, as additional pathways can activate the intracellular MAPK cascade that upregulates Pea3 transcription factor expression and activity (Fontanet et al., 2013; Haase et al., 2002; Helmbacher et al., 2003). To determine if the dorsal curvature in *3etv* mutants was due to a reduction in FGF signal transduction, we exposed larvae to the small molecule FGFR antagonist SU5402. Fish were treated with DMSO or SU5402 for 24-hour time periods between 18 hpf and 96 hpf and then we measured their body angle to see if there was a developmental window in which FGF signaling modulates axis curvature (Fig. 5D-T). SU5402 had no effect on body axis curvature when exposure started earlier than 24 hpf (Fig. 5D-F). By contrast, exposures starting between 24-60 hpf resulted in embryos with CTU phenotypes (Fig. 5G-R). Treatments with SU5402 starting at 72 hpf resulted in only slight dorsally curved tails. This curvature was specific to SU5402 as treatment with DMSO alone did not result in a CTU phenotype. These results suggest that FGF signaling has a role for modulating body axis straightening between 24 and 72 hpf and prevents excessive dorsal tail movement. As SU5402 can inhibit other pathways in addition to FGF, we further investigated if FGF was responsible for this phenotype by expressing a dominant-negative (dn) *fgfr1a* using the Tg(*hsp70l:dnfgfr1a-eGFP*), which expresses the dnFgfr1a following heat-shock (Lee et al., 2005). We heat-shocked embryos at 30 hpf and measured their tail angles at 72 hpf. Heat shocked non-transgenic wild-type fish had a straight body axis, whereas the *dnfgfr1a-eGFP* expressing fish had the CTU phenotype (Fig. 5S-U). All together our results argue strongly that FGF signaling normally acts to attenuate the body straightening mechanism once the body reaches a straight axis.

**Figure 5.**
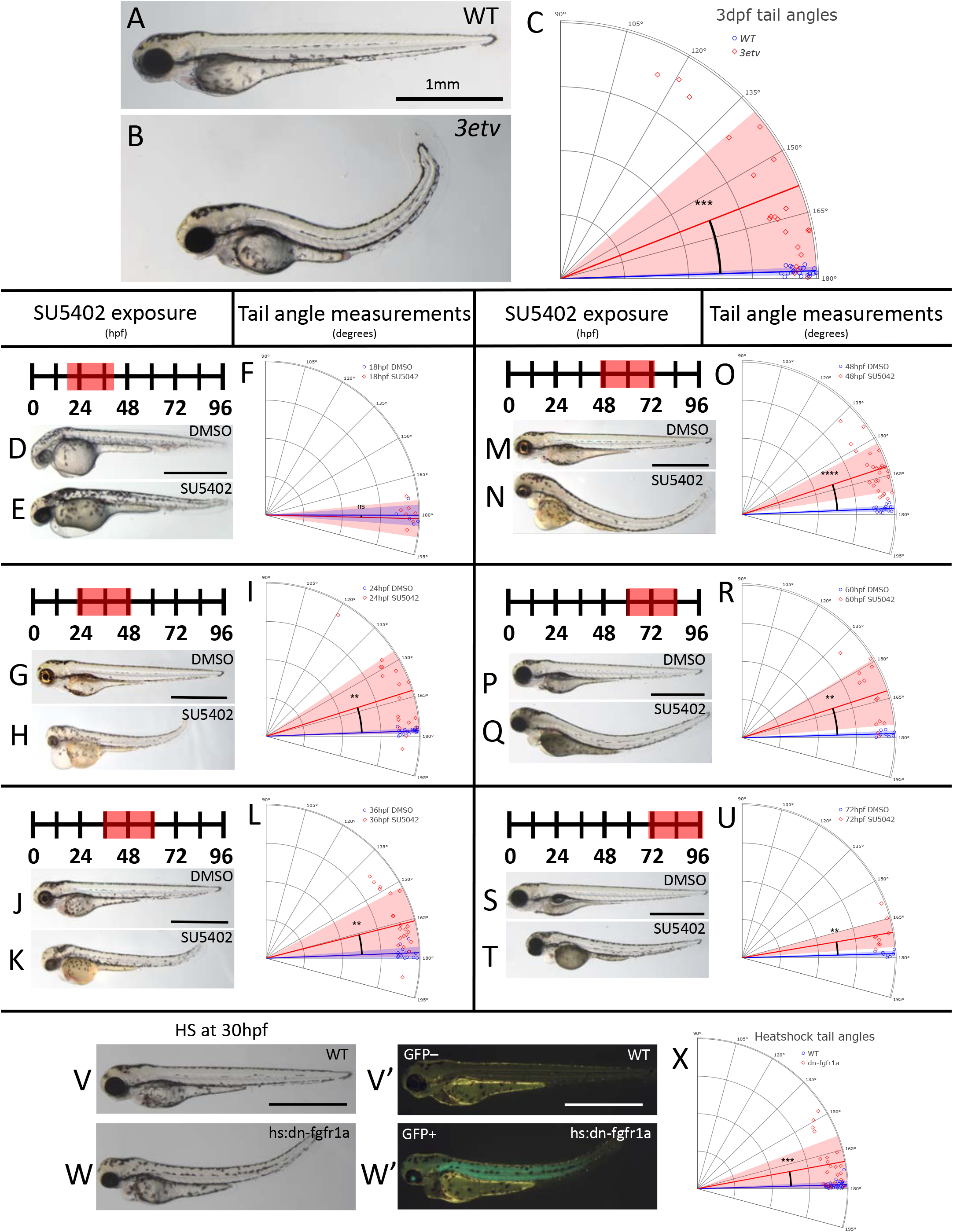
Fgf signaling modulates body axis straightening. (A-B) Light micrographs of wild-type and *3etv* 3 dpf larvae. Wild-type have a straight body axis (A) whereas *3etv* larvae have dorsally curved tails (B). (C) Polar plot of wild-type (blue diamond) and *3etv* (red circle) body axis angles. Solid lines show average of wild-type or *3etv* measurements, light blue and light red sectors show standard deviation. (D-U) Images of larvae following 24-hour long DMSO or SU5402 exposure with tail angle quantification following treatment. Light micrographs (V, W) and corresponding fluorescent micrographs (V’, W’) of 3 dpf larvae heat-shocked at 30 hpf. Wild-type heat-shocked fish have a straight body axis (V) and no GFP expression (V’) at 5 dpf. Fish with tg(hs:dn-fgfr1a-eGFP) have dorsally curved tails (W) and are GFP-positive (W’) following heat-shock. (X) Polar plot quantification of body curvature for heat-shocked wild-type and tg(hsp70l:dn-fgfr1a-GFP) larvae. All scale bars are 1mm.

### *3etv* mutants downregulate polycystin genes

While the CTU phenotype has not previously been observed in perturbations of FGF signaling, extreme dorsal tail curvature is consistently found in fish with loss of the polycystin genes *pkd1a, pkd1b*, and *pkd2* (Bisgrove et al., 2005; Mangos et al., 2010; Schottenfeld et al., 2007; Sternberg et al., 2018). The polycystin proteins localize to the membrane of immotile cilia where they complex to form non-selective Ca^2+^ permeable cation channels thought to serve as mechanosensors. Mutations in the polycystin genes are the primary cause of Autosomal Dominant Polycystic Kidney Disease (ADPKD) in humans, and in zebrafish polycystin mutations lead to defects in collagen deposition and cartilage formation, renal cysts, and abnormal LR axis patterning in addition to CTU. However, a link between polycystin function and FGF signaling has not been previously reported.

To identify if polycystin gene expression was altered in *3etv* mutants we first profiled polycystin gene expression in our 30 hpf RNA-seq data set and found that *pkd1a, pkd1b*, and *pkd2* were reduced in mutant embryos (Fig. S3). To confirm this finding and see if there were changes in polycystin expression during the tail straightening process we performed RT-qPCR on single embryos at different stages of larval development from 24-48 hpf to quantify the expression levels of polycystin genes in *3etv* mutants relative to wild-type (Fig. 6A). We found that polycystin gene expression was downregulated in the *3etv* mutants; *pkd1b* expression was reduced in *3etv* mutants at all stages tested, *pkd2* levels were lower at 24 and 36 hpf, and *pkd1a* was down at 36 hpf. To confirm the downregulation of polycystins and observe how expression was altered, we used RNA *in situ* hybridization to assay *pkd1b* expression in wild-type and *3etv* embryos at 26 hpf, a stage when *pkd1b* is normally expressed throughout the spinal cord and scored the embryos for either high or low expression. Consistent with our RT-qPCR analysis, we found evidence for reduced *pkd1b* expression in all *3etv* mutants (Fig. 6B-D).

**Figure 6.**
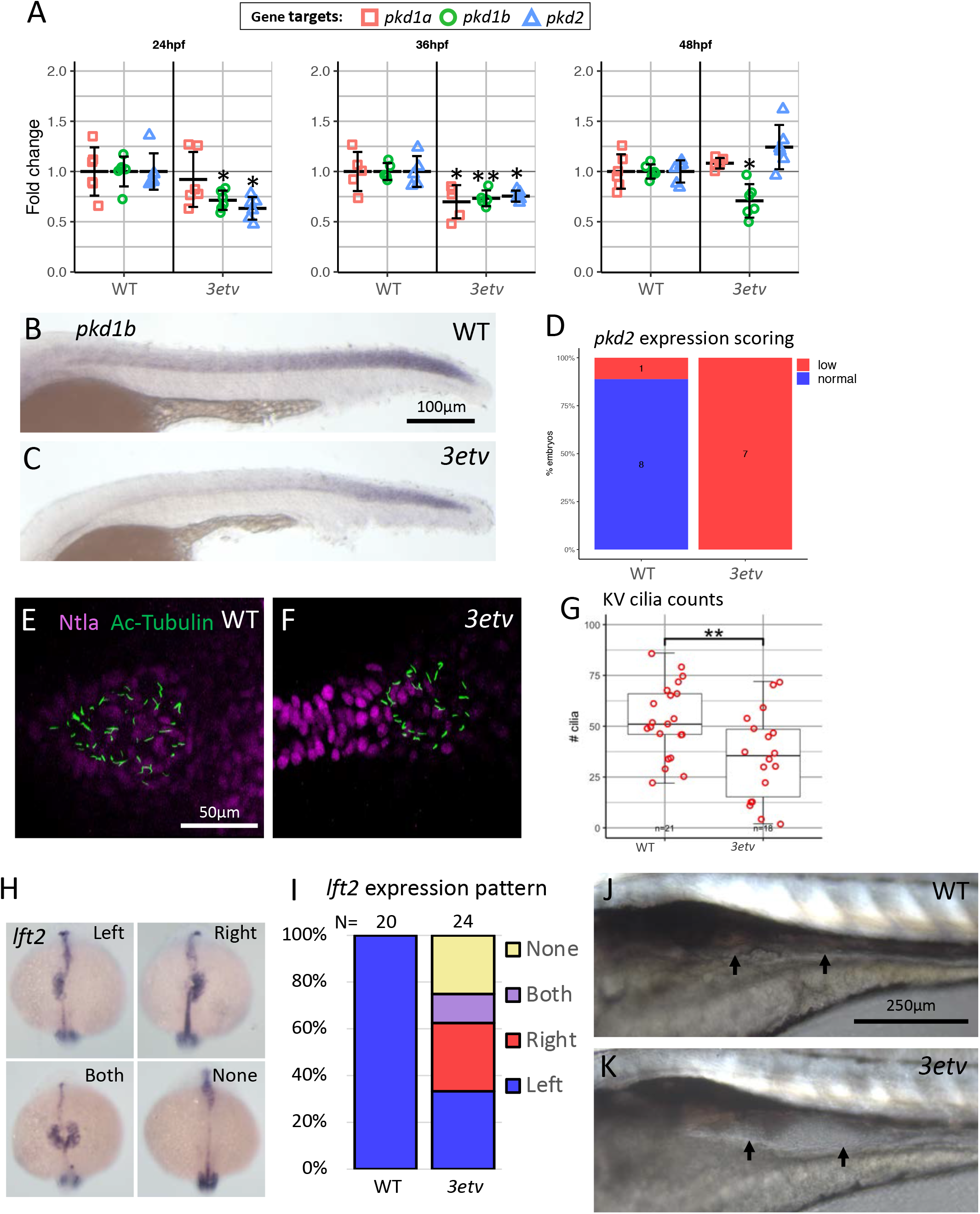
*3etv* mutants downregulate polycystin genes. (A) Fold change of polycystin gene expression at three time points from 24 hpf-48 hpf. Error bars are standard deviation. (B-C) ISH for *pkd1b* at 30 hpf. (B) Wild-type expression of *pkd1b* extends the length of the spinal cord neural tube, while *pkd1b* in *3etv* (C) mutants is reduced and restricted to the posterior neural tube. (D) Qualitative scoring of *pkd1b* ISH staining in wild-type and *3etv* mutants into low and high expression groups. (E-F) Max projections of Z-stacks imaging KV cilia in wild-type and *3etv* 10 ss embryos. (G) Boxplot of the number of cilia present in individual embryos; *3etv* mutants had fewer KV cilia than wild-type. (H) Expression patterns of *lft2* ISH staining observed in *3etv* 19 ss embryos. In addition to the normal left side expression, mutants displayed inversed right side expression, expression on both left and right sides, and no expression of *lft2*. (I) Stacked bar plot representing the percentage of wild-type and *3etv* embryos with the different *lft2* expression patterns. (J-K) Nemarski-DIC micrographs of wild-type and *3etv* trunks at 4 dpf. The pronephros is swollen in *3etv* mutants (arrows).

Given the downregulation of polycystin genes in *3etv* mutants, we next asked if *3etv* embryos have additional phenotypes that would be consistent with reductions of polycystin expression. For example, loss of Pkd2 function results in LR patterning defects and renal cyst (Obara et al., 2006; Sun et al., 2004). During normal vertebrate development, LR patterning is initiated by a structure located at the posterior end of the midline, Kupffer’s Vesicle (KV) or the node, respectively. Motile cilia present on the epithelial lining of these structures produce directed fluid flow that is thought to activate mechanosensing primary cilia on cells that are located on the left side of these structure, which in turn leads to the activation of asymmetric gene expression (Tabin and Vogan, 2003). We therefore stained for cilia located in the KV for wild-type and *3etv* embryos at 10 ss. We found that *3etv* mutants had significantly fewer cilia than WT on average, though the defect was variable (Fig. 6E-G). We also asked if *3etv* mutants result in defects in LR determination and stained embryos for LR asymmetry marker *lft2*, which is expressed on the left axis in the developing heart field. Expression was randomized in *3etv* mutants at 19 ss, with a mixture of correct left side expression, or incorrect right side, both sides, or no expression of *lft2* (Fig. 6H,I). The polycystin genes are required for normal kidney function and loss of polycystins results in the formation of kidney cysts. To determine if *3etv* mutants developed kidney cysts due to a reduction of polycystsin expression we imaged the 4 dpf larvae and found *3etv* mutants had a drastically enlarged pronephros, the zebrafish equivalent of cystic kidneys (Fig. 6J,K).

### Dorsal curvature in *3etv* mutants requires urotensin peptide function

The neural tube lumen has become a focus of research as a signaling hub that is involved in regulating tail curvature. In this model of body axis straightening, factors in the cerebrospinal fluid (CSF) transmit signals to trunk muscles via CSF-contacting neurons and neuromuscular junctions (Cantaut-Belarif et al., 2018; Sternberg et al., 2018; Zhang et al., 2018). Zhang et al. (2018) demonstrated that urotensin *urp1* and *urp2* neuropeptide ligands were necessary for body axis straightening to occur as loss of these neuropeptides resulted in a curly-tail down (CTD) phenotype. Conversely, overexpression of *urp1* is sufficient to cause the CTU phenotype, suggesting that tight regulation of urotensin neuropeptide expression and/or activity is critical for proper tail morphogenesis.

Since overexpression of urotensin peptides is sufficient to cause CTU we asked if the *urp1* and *urp2* genes were upregulated in *3etv* mutants. Our 30 hpf RNA-seq dataset suggested mutants had increased urotensin ligand expression (Fig. S3), and we validated this by measuring the relative levels of *urp1* and *urp2* mRNA in wild-type and *3etv* individual embryos using RT-qPCR between 24 and 36 hpf (Fig. 7A). We found that *urp1* and *urp2* were increased at 24 hpf, 30 hpf, and 36 hpf in *3etv* mutants. We next sought to answer if the increase of urotensin gene expression was due to an increase in the number of urotensin-expressing cells in *3etv* mutants or simply upregulation of urotensin. To this end we assayed *urp1* expression in wild-type and *3etv* mutants using RNA *in situ* hybridization at 30 hpf and counted the number of *urp1^+^* cells (Fig. 7B-D). We found that *3etv* mutant and wild-type embryos had comparable numbers of *urp1+* cells (Fig. 7D). However, the *3etv* mutants appeared to have increased *urp1* staining in the neural tube cells compared to wild-type, consistent with the increased *urp1* expression found by RT-qPCR. Based on these results it is possible that the *3etv* CTU phenotype results from increased urotensin gene expression and thus urotensin peptide production.

**Figure 7.**
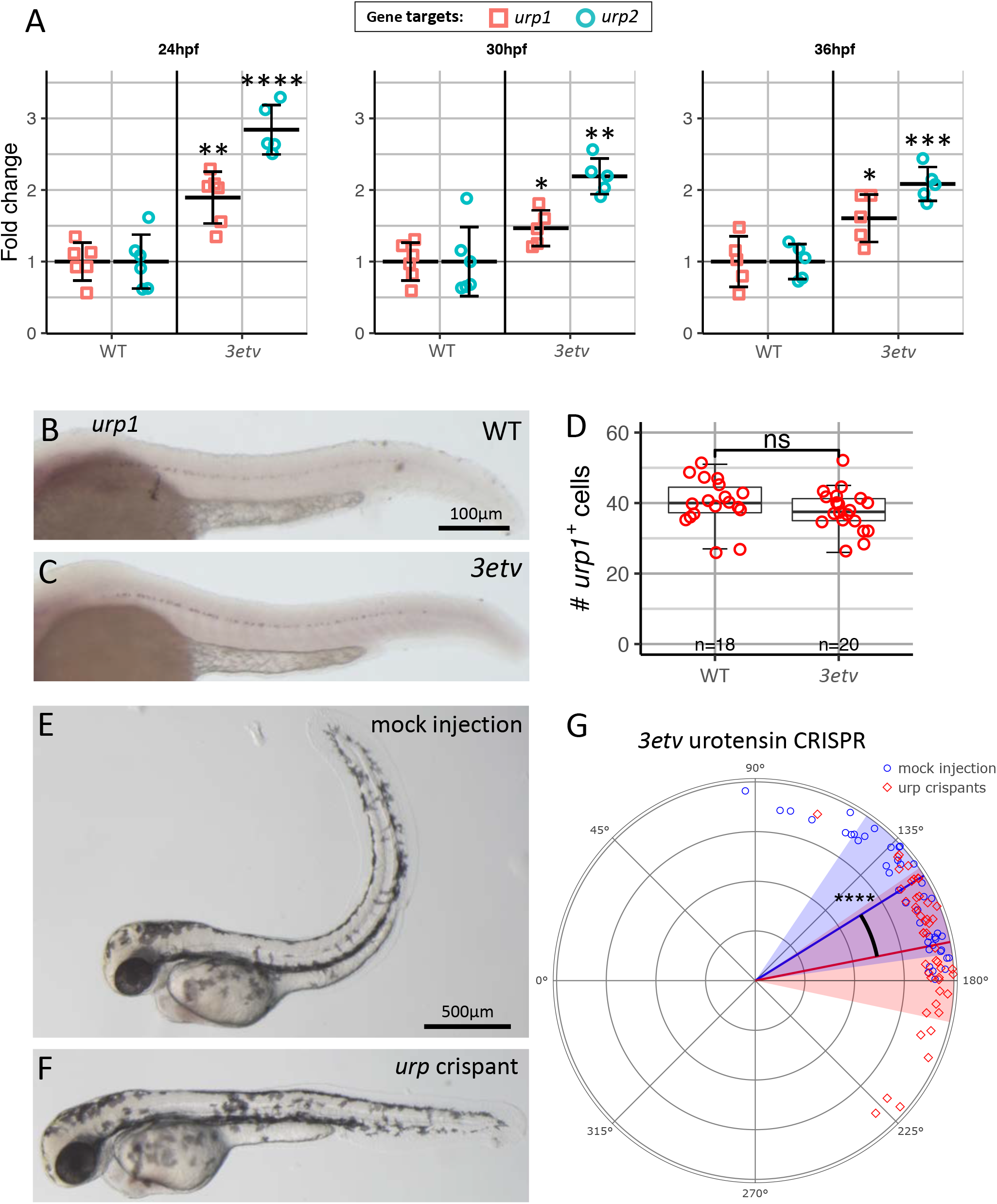
Dorsal curvature in *3etv* mutants is dependent on urotensin peptide genes. (A) Fold change of urotensin peptide gene expression at three time points from 24 hpf-36 hpf. Error bars are standard deviation. (B-C) ISH for *urp1* in 30 hpf wild-type and *3etv* larvae. (D) Boxplot of *urp1*-positive cell counts for individual fish; there was no difference between the number of *urp1*-positive cells in wild-type and *3etv* larvae. Fish were scored in a blinded manner and genotyped after counting was finished. (E-F) Bright-field micrographs of *3etv* mutants injected at the 1-2 cell stage with either no Cas9 mock injections (E) or urotensin targeted crispants (F). (G) Tail angle polar plot quantification of *3etv* mock injections and *3etv* urotensin crispants.

Finally, we asked if urotensin signaling was required for the *3etv* mutant CTU phenotype. We used CRISPR/Cas9 to induce indel mutations simultaneously in *urp1* and *urp2* for both wildtype and *3etv* mutant embryos. We first verified the efficacy of the *urp1* and *urp2* sgRNAs in wild-type embryos using HRMA and detected indels for both genes in the injected embryos (Fig. S4). Consistent with this, we found that injected embryos, hereafter referred to as urotensin crispants, developed a CTD phenotype at a frequency higher than mock injected controls (Fig. S4). We then injected embryos from a mutant incross and analyzed the tail angles of crispants at 2 dpf; *3etv* mutants were identified phenotypically by their otolith and fin defects. Mock injected *3etv* mutants had extreme dorsal tail curvature as previously observed; however, *3etv* urotensin crispants had significantly less tail curvature (Fig. 7E-G). These data show urotensin signaling is necessary for the CTU phenotype, and reduction of urotensin gene expression partially rescues dorsal tail curvature in *3etv* mutants. Together these data argue that FGF signaling via Etv4, Etv5a and Etv5b transcription factors are critical regulators proper tail morphogenesis.

## Discussion

Fibroblast growth factors are important and conserved regulators of vertebrate development. Though much is known about the developmental roles of individual ligands and receptors, much less is known about their downstream targets. In many developmental contexts FGF signaling activates the MAPK cascade, which in turn activates the Pea3 transcription factors. By focusing on the role of Pea3 genes we aimed to parse the broad effects of FGF by simplifying the scope to this single class of FGF-activated transcription factors. We used genetic mutations in the Pea3 transcription factors encoded by *etv4, etv5a*, and *etv5b* to better understand the biological aspects of FGF signaling they regulate and their function during early developmental events that could not be studied by previous MO knockdown studies.

### *etv4, etv5a* and *etv5b* act redundantly as positive mediators of FGF

The *etv4, etv5a*, and *etv5b* genes are well characterized downstream targets of FGF and consistent with this we found when all three genes are simultaneously mutated, embryos have phenotypes indicative of FGF signaling loss or reduction. This confirms these transcription factors are highly redundant and important positive regulators of the intracellular FGF response. However, in some cases the phenotypes in *3etv* mutants are not as severe as loss of particular FGF ligands or receptors. For example, *3etv* mutants have variable defects in pectoral fin development while mutations in either the FGF ligands encoded by *fgf24* or *fgf10*, or triple mutants for the FGF receptor encoding genes *fgfr1a;fgfr1b;fgfr2*, result in a complete absence of pectoral fin development. The fin defects observed in the *3etv* mutants suggest these factors are not required for the FGF-dependent initiation of the fin bud development, but instead play important roles during the fin outgrowth phase that is maintained by an FGF feed-forward loop. As another example, we found that the *3etv* mutants have a defect in MHB development that is less severe that loss of Fgf8a function. Specifically, *fgf8a* mutant embryos lack the midbrain-hindbrain junction, while *3etv* mutants do not. However, reduced expression of *pax2a* in the MHB of *3etv* mutants relative to wild-type embryos suggest that these three genes play a role in MHB development, albeit minor. While *3etv* mutants have phenotype suggesting partial loss of FGF signaling in some cases, the otic vesicle and LR patterning defects exhibited by *3etv* mutants are more akin to a complete loss of Fgf8a. Specifically, similar to *fgf8a* mutants, *3etv* mutants have reduced expression of *pax2a* in otic placode, have significantly smaller otic vesicles than wild-type, and form only one instead of two otoliths. With regard to LR patterning, *3etv* triple mutants have similar LR asymmetry defects to those reported for *fgf8a* mutants (Albertson and Yelick, 2005; Neugebauer et al., 2009). Together these results suggest that loss of *etv4, etv5a* and *etv5b* does not have uniform effects on FGF signaling, even in developmental events that are regulated by the same FGF ligand, such as Fgf8a.

The functions of *etv4, etv5* and *etv5b* during zebrafish embryogenesis have previously been investigated using MOs (Znosko et al., 2010). This study found that triple knockdowns resulted in phenotypes similar to our *3etv* mutants, though in some cases the morphants have more severe early developmental phenotypes than we observe. In particular, while both our study and that of Zonosko et al. (2010) found defects in MHB development, triple MO KD embryos completely lacked the MHB junction, similar to *fgf8a* mutants, while *3etv* mutants had only mild defects in the expression of *pax2a* but formed an apparently normal MHB. We do not know why MO knockdown cause a more severe effect on MHB development than genetic mutants, though there are several possible explanations. First, all three of our mutated genes are predicted to produce transcripts with premature stop codon and, consistent with this, our RT-qPCR analysis indicates that all three mutant transcripts appear to be subject to nonsense-mediated decay (NMD). Though our RT-qPCR analysis revealed minor evidence for genetic compensation in the single mutants (e.g. *etv5a* transcription is increased roughly 1.5 fold in *etv4* mutants), it is not likely genetic compensation could explain the differences we see between triple MO knockdown and the triple genetic mutants.

A more likely reason why genetic mutants may have a less severe phenotype than MO knockdown embryos is that mRNA for the *etv4, etv5a*, and *etv5b* genes are maternally provided, so differences in phenotypes could be due to varying levels of maternal contribution between the two experimental paradigms. For example, Znosko and colleagues used translation-blocking MO that can block the production of proteins from both maternal and zygotic mRNAs and in theory could prevent the production of 3Etv proteins during early embryogenesis (Znosko et al., 2010). In contrast, our genetic mutants removed maternal contribution of at most two of Etv4/5 genes. In generating the *3etv* mutants, we routinely incrossed parents that are homozygous mutant for two of the three genes and heterozygous for the third, for example *etv4^-/-^;etv5a^-/-^;etv5b^+/-^*. Thus, these embryos would have no maternal contribution for two of the three genes, *etv4* and *etv5a*, and only half the maternal contribution from the third gene, *etv5b*. It is therefore possible that this reduced level of maternal product is still sufficient to allow near normal midbrain-hindbrain development while triple MO knockdown embryos do not produce a MHB. However, we have not found any differences in the phenotypes of *3etv* mutants generated from these crosses and those generated from crosses between triple heterozygous parents arguing that levels of maternal gene product may have minimal effect on phenotypic outcome. Regardless, because the indel mutants survive to 13 dpf, we have been able to more fully assess the roles of *etv4, etv5a* and *etv5b* during both embryogenesis and early larval development- a timepoint inaccessible to functional studies with MO.

### Etv4, Etv5a and Etv5b contribute to normal development of posterior mesoderm

The role for FGF signaling in posterior mesoderm formation during gastrulation has been well established in vertebrates and the expression of the Pea3 genes found in presomitic mesoderm and newly formed somites is dependent on FGF signaling (Nguyen-Chi et al., 2012). Combinatorial loss of certain FGF receptors or ligands, or chemical inhibition of FGF signal transduction during gastrulation results in either a reduction or complete absence of posterior mesoderm formation. For example, *fgf8a^-/-^;fgf24^MO^* morpho-mutants, or embryos expressing a dominant-negative Fgfr1a receptor, have greatly reduced or absent production of posterior mesoderm (Draper et al., 2003; Griffin et al., 1995). In contrast, we found that *3etv* mutants produce nearly normal amounts of posterior mesoderm, but also found they have significant defects in gene expression that appear to lead to somite patterning defects. For example, previous studies using chemical inhibition have found evidence that FGF signaling plays a role in the development of somite boundaries and transient inhibition during somitogenesis can prevent boundary formation that then results in somites twice the normal width (Sawada et al., 2001). Indeed, we found similar defects in somite boundary formation in our *3etv* mutants that in some cases lead to the fusion of adjacent somites, a phenotype identical to that caused by blocking FGF signaling. This fused somite phenotype is also similar to phenotypes observed in *ripply1, ripply2*, and *tbx6* mutants (Ban et al., 2019; Kinoshita et al., 2018; Windner et al., 2015), suggesting that FGF is acting through *etv4, etv5a*, and *etv5b* to regulate the expression these genes. Consistent with this, we found that expression of *ripply1, ripply2* and *tbx6* are down regulated in *3etv* mutants, providing a genetic link between 3Etv function and somite boundary formation.

### FGF signaling regulates straightening of the posterior body axis after somitogenesis

Perhaps the most surprising finding from our study is that the 3Etv transcription factors play a necessary role in regulating body axis straightening, a process not previously known to involve FGF signaling. During normal development the trunk and tail initially form with a ventral-ward curvature that then straightens until roughly 30 hpf when a straight horizontal body axis is achieved, and defects in this process are linked to idiopathic scoliosis (Cantaut-Belarif et al., 2018; Grimes et al., 2016; Thouvenin et al., 2020). The straightening process appears to be driven by contractions of muscles located in the dorsal half of each somite that are triggered by the urotensin neuropeptides, Urp1 and Urp2, in the cerebrospinal fluid (CSF) within the spinal cord central canal (Zhang et al., 2018). *urp1* and *urp2* are expressed in and secreted from CSF-contacting neurons which line the brain ventricles and spinal cord central canal, and Urp1 and Urp2 are then thought to activate their receptor, Uts2r3, which is specifically expressed by dorsally located slow-twitch muscles. Morpholino knockdown of *urp1* and *urp2* prevents tail straightening and results in embryo with CTD phenotypes. Similarly, mutations in several genes that encode proteins required for motile cilia function, like *zmynd10*, which is necessary for assembly of axonemal dynein arms (Zariwala et al., 2013), result in embryos with CTD. These results have led to the proposal that activation of dorsal muscle contraction requires the urotensin peptides be distributed within the CSF by the motile cilia lining the spinal cord central canal.

In contrast, mutations in the polycystic kidney disease genes *pkd1 and pkd2* result in embryos with curly-tail up phenotypes (CTU), similar to *3etv* mutants. *pkd1* and *pkd2* encode protein components of a Ca^2+^-activated nonspecific cation channels that localizes to cilia and is thought to function in cilia-based mechanosensation. As such, *etv4, etv5a* and *etv5b* appear to function in a pathway with the polycystin proteins to attenuate the urotensin-promoted muscle contractions once the tail has fully straightened, thereby preventing the tails from continuing to curl upwards. Thus, correct body axis straightening appears to result from coordinated muscle contraction-promoting and contraction-attenuating signals.

While it was possible that the role of the 3Etv transcription factors in body axis straightening was independent of FGF signaling, we showed that blocking FGF receptor function between 24 and 72 hpf with either chemical inhibition of ectopic expression of dnFgfr1a resulted in a CTU phenotype identical to *3etv* mutants. From these results we conclude that FGF signaling plays a necessary role in regulating body axis straightening during embryogenesis. There are several possible explanations for why FGF signaling has so far not been implicated in axis straightening. First, it is possible that redundant FGF ligands are involved so single ligand knockouts would not display this phenotype. Redundancy of FGF ligands is well established in several developmental contexts in zebrafish (e.g. Draper et al., 2003; Manfroid et al., 2007). Second, FGF is required throughout posterior mesoderm development so drastic reduction of FGF signaling prior to body axis straightening would likely be masked by severe defects in the tail, if posterior mesoderm formed at all.

Current studies have identified two opposing pathways regulating body axis orientation, one, urotensin, that promotes dorsal-ward curvature and a second, polycystin, that appears to attenuate the dorsal-ward curvature once a straight tail is achieved (Schottenfeld et al., 2007; Zhang et al., 2018). Our results indicate that reduced FGF signaling, as we argue occurs in the *3etv* mutants, affects the expression of genes in both pathways. Specifically, we found that the straightening-promoting urotensin genes were upregulated, while the straightening-attenuating polycystin genes were downregulated in *3etv* mutants. Altered gene expression in any one of these pathways alone could result in the CTU phenotype, but it is also possible the phenotype is due to their additive effects. Our results blocking FGF signaling with SU5402 shows that maximum tail curvature defects occur when signaling is blocked between 24-72 hpf, a timepoint after the tails of normal embryos have completed straightening. This result argues that FGF is required not only for regulating correct axis orientation at 30 hpf, but also for continuously maintaining a straight body through 72 hpf. Urotensin expression is gradually reduced in spinal cord neurons after 30 hpf in wild-type embryos (Quan et al., 2015); however loss of FGF activity after as late as 72 hpf is still sufficient for the progression of CTU. Therefore, mis-regulation of polycystins may be the primary cause of the CTU phenotype in *3etv* mutants rather than upregulation of urotensin signaling.

### *3etv* and polycystin may have linked role in cilia function

In addition to tail straightening defects, the polycystin mutants have LR patterning defects and, as their name implies, form kidney cyst – two phenotypes we have also identified in *3etv* mutants (Bisgrove et al., 2005; Schottenfeld et al., 2007; Sun et al., 2004). These shared phenotypes suggest these genes function in a similar pathway. Interestingly these phenotypes have also been reported to result from global reduction of FGF signaling during early development in zebrafish (Neugebauer et al., 2009). FGF signaling, in combination with *pkd2*, is required for specification of the flow sensing cells within the LR organizing center in *Xenopus* (Schneider et al., 2019). However, that study concluded the FGF activity was mediated by the Ca^2+^ branch of FGF signal mediation and *pkd2* was in a separate parallel pathway as they found no interaction between FGF and *pkd2*. Conversely, our research suggests a major role for FGF to directly control the expression of the polycystin genes via the 3Etv transcription factors. A direct link between polycystin protein function and cilia function is well established, and it has also been proposed that FGF signaling regulates cilia formation (Liu et al., 2011; Neugebauer et al., 2009; Riley et al., 1997). Furthermore, the phenotypes shared by both *3etv* and polycystin mutants can also result from other mutations that affect cilia formation and/or function, so called ciliopathy phenotypes (reviewed in Reiter and Leroux, 2017). Thus, similar phenotypes may arise from independent effects of polycystin and FGF pathways on cilia formation and function.

In addition to the polycystin-associated phenotypes, *3etv* mutants have another phenotype that is also linked to defects in ciliogenesis, lending further support to the idea that FGF signaling, through 3Etv transcription factors, plays widespread roles in ciliogenesis. Both *3etv* and *fgf8a* mutants produce otic vesicles that are smaller than wild-type embryos and that have only a single otolith instead of two. During normal development, otoliths, which are composed of calcium carbonate and function as part of the auditory and vestibular systems, are formed by the localized accretion of smaller granules that are floating within the otic vesicle. Riley and colleagues showed that the accretion that forms the otoliths occurred at the tip of specialized cilia, called tethering cilia, located in the anterior and posterior regions of the otic vesicle and that removing the cilia blocked otolith formation (Riley et al., 1997). Fgfr1a morphants have severely reduced tethering cilia fail to form the anterior otolith, consistent with *fgf8a* and *3etv* mutants (Neugebauer et al., 2009; Scholpp and Brand, 2004). However, it remains to be seen if other defects observed in *3etv* mutants, such as in fin and mesoderm development, can also be directly linked to defects in cilia development or function.

In conclusion, the *3etv* genes represent just one target for one of the several intracellular pathways activated by FGF, thus allowing us to parse FGF signaling at a finer resolution than possible in broader losses of FGF signaling that occurs in receptor and ligand mutants or via chemical inhibition. Our results strongly argue that one important responses downstream of the FGF/3Etv pathway is to regulate cilia formation and/or function.

## Methods

### Husbandry

The wild-type strain NHGRI-1 was used for the generation of *etv4^uc77^, etv5a^uc80^*, and *etv5b^uc82^*. Mutant fish were outcrossed to wild-type strain AB, and wild-type controls were used in experiments. Zebrafish staging and husbandry was performed as previously described (Westerfield, 2007).

### CRISPR/Cas9 injection and mutant allele generation

A single-stranded guide RNA (sgRNA) and Cas9 mRNA were prepared as described previously (Leerberg et al., 2019; Moreno-Mateos et al., 2015). Sequences for all sgRNA guide oligos and scaffold oligo are listed in Table S1. An injection solution was prepared with 12ng/μL of guide RNA, 30ng/μL Cas9 mRNA, 300mM KCl, and 0.050% phenol red. 2nL of injection solution was injected into the yolk of 1-cell stage zebrafish embryos. Cutting at the targeted locus was confirmed by High Resolution Melt Analysis (HRMA) at 24 hpf with 8 injected embryos and 8 uninjected controls (Dahlem et al., 2012). Sperm from injected males was screened by HRMA for germline transmission of alleles with indels. Genomic DNA from injected male founders with frameshift-causing indels were sequenced with Sanger sequencing; frameshift mutant alleles were confirmed by sequencing cDNA from outcrossed embryos.

The sgRNAs for urotensin crispants were generated as described above. We designed 2 sgRNAs for each *urp1* and *urp2* and injected all 4 simultaneously to ensure knockdown of urotensin expression. Crispant injections consisted of 17.5ng/μL of each urotensin sgRNA, 75ng/μL Cas9 mRNA, 300mM KCl, and 0.050% phenol red. The control mock injections did not contain Cas9 mRNA. 2nL of either the mock or urotensin CRISPR/Cas9 mix was injected into the yolk just beneath the cell in 1-2 cell stage embryos from an etv4^+/-^;etv5a^-/-^;etv5b^-/-^ incross. *3etv* mutants were identified at 2 dpf and imaged for tail angle quantification. We then performed HRMA on 16 mock and crispant embryos to confirm indel formation at the urotensin sgRNA target sites.

### Genotyping

Mutants for *etv4, etv5a*, and etv5b were genotyped using standard PCR conditions for Taq polymerase using genomic DNA extracted from caudal tail samples in lysis buffer with proteinase K digestion (Westerfield, 2007). For the RT-qPCR and RNA-seq experiments, *3etv* embryos were identified by visible mutant phenotypes prior to RNA extraction. Primer oligo sequences are listed in Table S1.

### RT-qPCR

RNA isolation and cDNA generation were performed as detailed in (Leerberg et al., 2019). For the compensation experiment, embryos from a heterozygous incross had tail biopsies taken and placed in DNA lysis buffer for genotyping while the rest of the embryo was snap frozen in liquid nitrogen and stored at −80°C until homozygous mutants were identified by genotyping. For the polycystin and urotensin fold-change quantification experiments, mutants were identified by visually phenotyping at the respective timepoints.

RT-qPCR was performed on cDNA generated from individual embryos. Each group was composed of 5-6 single embryo biological replicates and 3 technical replicates per gene target. Fold-change expression values were calculated in the Bio-Rad CFX Maestro software. RT-qPCR plots were generated using the R package ggplot2 and contain the fold-change values of all biological replicates plotted along with mean fold-change for each group and standard deviation error bars (Wickham, 2011). Fold-change values are normalized to wild-type expression for each gene target. Primer oligo sequences are listed in Table S1.

### Cartilage staining

Larvae were fixed in 4% paraformaldehyde (PFA) overnight at room temperature and cartilage staining with Alcian blue was performed as previously described (Walker and Kimmel, 2007). The tissue treated with 1% Trypsin dissolved in a saturated tetraborate solution to clear tissue. Viscerocranial and neurocranial cartilage were dissected with fine tipped needles. Cartilage was flat mounted in 75% glycerol for imaging on a dissecting scope.

### RNA *in situ* hybridization

Samples were fixed in 4% PFA overnight at 4°C. Samples were dehydrated with 100% methanol and stored at −20° for at least 24 hr. Embryos >30 hpf were bleached for ~10 min prior to proteinase K digestion in 3% H_2_O_2_, 0.5% KOH. Color *in situ* hybridizations were performed by a procedure similar to that of (Thisse and Thisse, 2008) with the exception that 5% dextran sulfate was included in the hybridization solution. RNA probes that detect the following genes were used: *tbxta* (Schulte-Merker et al., 1992); *myod1* (Begemann and Ingham, 2000); *pax2a* (Krauss et al., 1991); and *etv4* (Münchberg et al., 1999). Probes for *dlc, lft2, pkd1b, urp1*, and *xirp2a* antisense probes were generated from PCR amplified products from cDNA using a T7 tagged reverse primer; probe primer sequences are listed in Table S2. For experiments where stained embryos were scored, wild-type and *3etv* embryos were processed together in the same tube for ISH staining so they could be imaged and scored in a blinded manner, then genotyped only after scoring was finished.

### RNA-seq data analysis

RNA isolation on single embryos was performed as detailed in (Leerberg et al., 2019). mRNA-seq libraries were generated using standard Illumina protocols for poly-A enriched, strand-specific libraries. PE150 reads were generated with the NovaSeqS4 sequencing platform, mapped to the GRCz11 genome using the STAR aligner (2.5.3a), and gene hit count matrices were generated with HTSeq-count (0.6.1p1) (Anders et al., 2015; Dobin et al., 2013). DESeq2 with apeglm effect size calculation was used R for DGE data analysis (Love et al., 2014; R Core Team, 2018; Zhu et al., 2019). For data visualization, heatmaps were generated with Heatmap2 in the gplots package with the RColorBrewer “RdBu” color pallete and volcano plots were generated with EnhancedVolcano (Blighe, 2019; Neuwirth, 2014; Warnes et al., 2020). For gene ontology (GO), differentially expressed genes that were downregulated in *3etv* mutants were submitted to DAVID for functional annotation clustering using GOTERM_DIRECT_BP (Huang et al., 2008; Huang et al., 2009). Enriched GO terms were processed with REVIGO and visualized using CirGO (Kuznetsova et al., 2019; Supek et al., 2011).

### Body axis and somite curvature analysis

Body axis and somite curvature was quantified for zebrafish larvae in ImageJ using the Angle tool. Somite curvature was measured as the angle formed where the dorsal somite boundary intersected with the notochord. The body curvature angle was found by taking the angle formed by the centroid of the otic vesicle, the center of notochord dorsal to the anal vent, and the tip of the tail. For fish with body curvature greater than 180°, in the case of curled down fish, the body angle was calculated by subtracting the angle measured in ImageJ from 360°. A visualization of how the body axis and somite curvature angles were measured for zebrafish larvae can be found in Figure S5.

### SU5402 exposure

Groups of 6-8 larvae were exposed to SU5402 (Selleck Chemical Llc, Cat. # S7667) in 24 well tissue culture dishes (Corning, Cat. # 353226). For each exposure timepoint effective doses of SU5402 were found by exposing groups of larvae to each concentration in a 2x dilution series from 0.625-10mM SU5402 and a DMSO control for 24 hours. The concentration of DMSO was kept constant in all exposures at 0.1%. Larvae were assessed for survival and imaged after immediately after the 24-hour exposure. The highest concentration at which the fish remained viable following SU5402 exposure was determined as the effective dose. For replication, exposures were performed a minimum of 2 times at the effective dose.

### dnfgfr1a-eGFP heat-shock

dnfgfr1a-eGFP hemizygous fish were outcrossed with AB wild-type fish. At 30 hpf individual larvae were placed into PCR plate wells with 100μL E3 embryo water and heat-shocked in a PCR thermocycler at 38°C for 30 minutes. Heat-shocked larvae were screened at 72 hpf for GFP fluorescence and imaged for body curvature measurements.

### Antibody staining

Samples were fixed and prepared the same as ISH samples. Following fixation, immunohistochemistry was performed as previously described (Leerberg et al., 2017). Rabbit-anti-Ntla antibody (Schulte-Merker et al., 1992) was used at a dilution of 1:5000 in PBS-triton, and mouse-anti-Ac-Tubulin antibody 6-11B-1 (Millipore-Sigma, Cat. # MABT868) was used at 1:100 diluted in PBS-triton. Antibodies were detected using Alexa Fluor secondary antibodies 488 goat-anti-mouse-IgG and 594 goat-anti-rabbit-IgG (Thermo Fischer Scientific, Cat. #s A-11001 and A-11012). Embryos were flat mounted in 75% glycerol, and heads were removed for DNA extraction and genotyping. Embryos were imaged using an Olympus FV1000 laser scanning confocal microscope.

### Statistics

Statistics were calculated using a 2-tailed Welch’s T-Test for samples with unequal variance. In experiments where multiple hypotheses were tested, p-values were adjusted with false discovery rate (FDR) correction. Significance values key: * p < 0.05, ** p < 0.01, *** p < 0.001, **** p < 0.0001.

## Acknowledgements

Tg(*hspl70*:DN-*fgfr1a*-eGFP) fish were a generous gift from Dr. Sharon L. Amacher at Ohio State University. The *lft2* probe was a generous gift from Dr. Debora Yelon at University of California, San Diego. We thank Celina Juliano for critical comments on the manuscript. This work was supported by the National Institute of Child Health and Development [1R01HD081551 to B.W.D.] and by the National Institute of General Medical Sciences [T32GM007377 to M.E.M.] The authors declare no competing interests.

## Data Availability

10 ss (<u>GSE155013</u>) and 30 hpf (<u>GSE155014</u>) RNA-seq datasets are available at the NCBI Gene Expression Omnibus (GEO) repository.

## Supplementary information

Figure S1. **Mutant Pea3 gene allele sequences** Alignments for wild-type (top) and Pea3 mutant (bottom) gDNA/cDNA sequences along with a description of the indel. Alignments are numbered from the start of the specified exon.

Figure S2. **RNA-seq DGE and quality assessment** (A) PCA plot of mRNA sequencing datasets for wild-type and *3etv* embryos at 10 ss and 30 hpf; the three biological replicates for each group cluster according to stage and genotype. (B-C) Volcano plots depicting significantly differentially expressed genes at 10 ss and 30 hpf timepoints (p-value ≤0.01 and fold change ≥1.5).

Figure S3. **Expression of polycystin and urotensin genes in the 30 hpf RNA-seq dataset** (A) Heatmap of polycystin genes and urotensin genes expression Z-scores in 30 hpf wild-type and *3etv* mutants. Blue color represents expression below the mean expression of the row and red is higher expression than the row mean. Genes with significant differential expression between wild-type and *3etv* mutants are annotated with asterisks.

Figure S4. **Urotensin sgRNA cutting validation** (A-B) High resolution melt analysis (HRMA) of 24 hpf CRISPR/Cas9 guides for *urp1* and *urp2*. 8 sgRNA only injected controls and 16 urotensin crispants were analyzed for each gene. Melt curves of amplicons for both genes all grouped together in control embryos, whereas melt curves were altered in the majority of urotensin crispants. (C-E) Urotensin crispants in the AB background had a low prevalence of CTD at 48 hpf. (F) Quantification of CTD phenotype in 48 hpf AB embryos injected with either sgRNA only control or urotensin CRISPR/Cas9 reagents.

Figure S5. **Tail angle measurement schematic** (A-B) Body curvature was determined by measuring the angle formed by points at the ear, the center of notochord dorsal to the anal vent, and the tip of the tail. Representative wild-type (A) and *3etv* (B) larvae drawings are overlaid on polar coordinate charts to depict how the curvature angle, Θ, is determined with these points on the body.

Table S1. **CRISPR/Cas9 sgRNA oligo sequences**

Table S2. **Primer oligo sequences**

Table S3. **10 ss RNA-seq DEGs**

Table S4. **30 hpf RNA-seq DEGs**

